# Transmission of pre-XDR and XDR-TB in the Mumbai Metropolitan Region, India

**DOI:** 10.1101/2021.02.02.429364

**Authors:** Viola Dreyer, Ayan Mandal, Prachi Dev, Matthias Merker, Ivan Barilar, Christian Utpatel, Kayzad Nilgiriwala, Camilla Rodrigues, Derrick W. Crook, the CRyPTIC Consortium, Nerges Mistry, Stefan Niemann

## Abstract

Multidrug-resistant (MDR) and extensively drug resistant (XDR) *Mycobacterium tuberculosis* complex (MTBC) strains are a great challenge for tuberculosis (TB) control in India. Still, factors driving the MDR/XDR epidemic in India are not well defined.

To address this, whole genome sequencing (WGS) data from 1 852 MTBC strains obtained from patients from a tertiary care hospital laboratory in Mumbai were used for phylogenetic strain classification, resistance prediction, and cluster analysis (12 allele distance threshold). Factors associated with pre-XDR/XDR-TB were defined by odds ratios and a multivariate logistic regression model.

Overall, 1 017 MTBC strains were MDR, out of which 57.8 % (n=591) were pre-XDR, and 17.9 % (n=183) were XDR. Lineage 2 (L2) strains represented 41.7 % of the MDR, 77.2 % of the pre-XDR, and 86.3 % of the XDR strains, and were significantly associated with pre-XDR/XDR-TB (P < 0.001). Cluster rates were high among MDR (78 %) and pre-XDR/XDR (85 %) strains with three dominant L2 strain clusters (Cl 1-3) representing half of the pre-XDR and two thirds of the XDR-TB cases. Cl 1 strains accounted for 52.5 % of the XDR MTBC strains. Transmission could be confirmed by identical mutation patterns of particular pre-XDR/XDR strains.

As a conclusion high rates of pre-XDR/XDR strains among MDR-TB patients require rapid changes in treatment and control strategies. Transmission of particular pre-XDR/XDR L2 strains is the main driver of the pre-XDR/XDR-TB epidemic. Accordingly, control of the epidemic in the region requires measures with stopping transmission especially of pre-XDR/XDR L2 strains.

## Introduction

Multidrug-resistant (MDR) tuberculosis (TB) caused by *Mycobacterium tuberculosis* complex (MTBC) strains resistant to at least isoniazid (INH) and rifampicin (RMP) poses a great challenge to global TB control. More than 400 000 new MDR-TB cases are notified annually ^1^; 50 % of these coming from India (27 %), China (14 %) and countries of the Russian Federation (9 %). This makes them the epi-centers of the current MDR-TB epidemic and key countries for the successful future intervention against MDR-TB^1^. Approx. 6 % of MDR-TB cases are already estimated to be extensively drug resistant (XDR, additional resistance to one of the fluoroquinolones [FQs] as well as to one of the injectable drugs, amikacin [AMI], capreomycin [CAP], or kanamycin [KAN])^2,3^.

The treatment of MDR-TB patients is longer, based on less effective, more toxic drugs, and in 2019, the cure rate was only 57 % on a global level^1,3^. With 39 %, the treatment success rate is even lower for XDR-TB patients^2^. As ineffective treatment is an important factor driving transmission^4^, the potential of MDR/XDR MTBC strains to transmit may be even higher compared to susceptible MTBC strains^5^.

As outlined above, India is particularly affected and, thereby becomes one of the main drivers of the current MDR/XDR-TB epidemic. Few studies on MDR/XDR-TB in India underlines the urgent need to gain more and better knowledge of the genetics of resistance determinants, evolution and transmission, of MDR/XDR MTBC strains in the region^6,7^. Indeed, it is of particular importance to understand the origins and driving forces of the MDR/XDR-TB epidemic in the country including ongoing transmission of already highly resistant clones^8–10^, but only few studies have used state-of-the-art whole genome sequencing (WGS) combined with epidemiological techniques to investigate transmission in India so far^11–14^.

To address these knowledge gaps, we performed a retrospective genomic epidemiological analysis based on WGS of 2 040 MTBC strains mainly from the Mumbai metropolitan region, India. The strains were obtained from a tertiary care hospital laboratory in Mumbai, which providing comprehensive drug susceptibility testing (DST) of MTBC strains. WGS data were used to determine MTBC-lineage, resistance to first- and second-line drugs, and transmission inference of MTBC strains based on sequence-based cluster analysis.

## Results

### Study population

Over the study period, MTBC strains were obtained from 2 040 patients. These strains were sent to the laboratory of the tertiary care hospital in Mumbai from private physicians practicing in India, mainly the Mumbai Metropolitan Region (Figure S1).

WGS data quality was sufficient for 1 852 strains (90.8 %), while WGS data of 188 isolates were excluded from data analysis: 22 had a coverage of below 40x, in seven samples the proportion of unambiguous reads were below 85 %, another 40 samples had mixed infections with two MTBC strains and 119 had technical errors arising due to discrepancies in levofloxacin drug susceptibility testing between genotypic and phenotypic DST (Figure 1).

**Figure 1.**
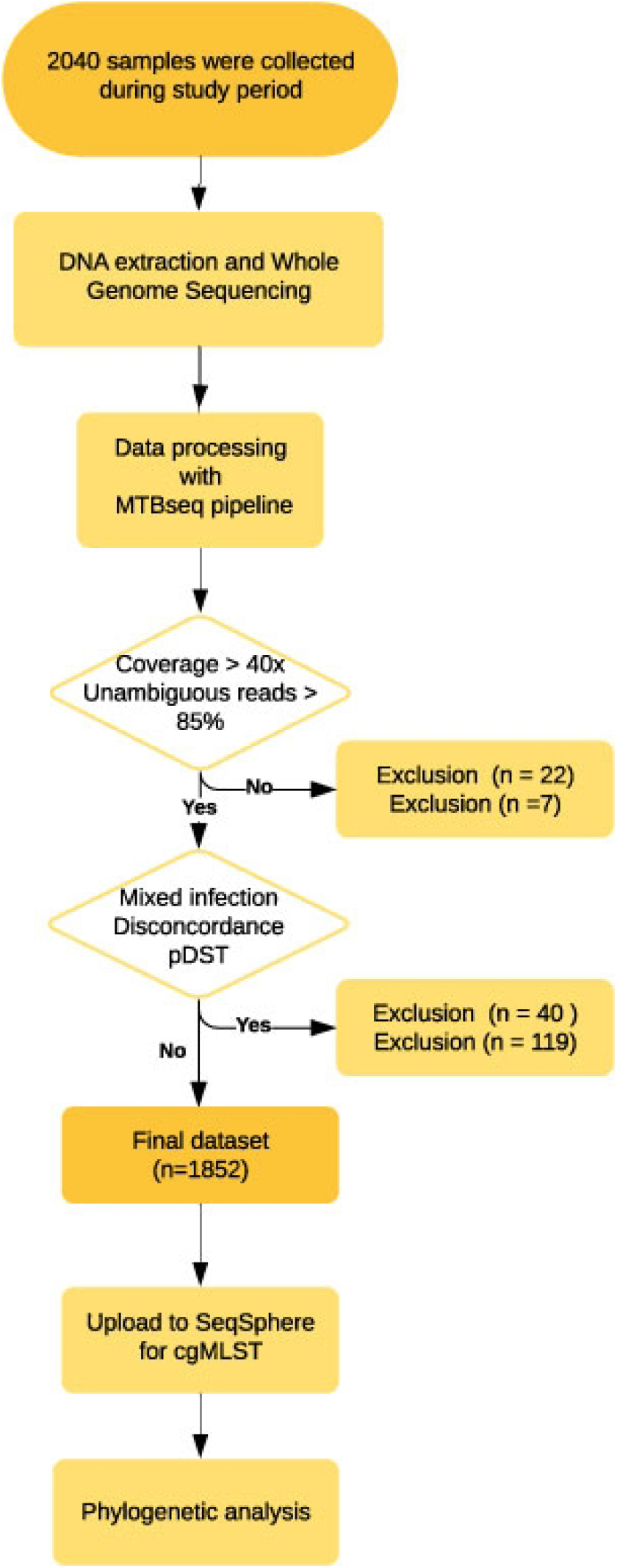
Study flowchart. In- and exclusion criteria for strains are reported in the two rhombuses. Final dataset consists of 1 852 strains.

Of the 1 852 patients, 55.5 % (n = 1 027) were female, and 44.5 % (n = 824) were male. TB cases were mostly diagnosed within the age group of 18 to 40 years (n = 1 081; 58.4 %) and were widely distributed in Mumbai and its suburban region (Figure S1, Table S3). The mean age of the whole population was 33.7 years (SD 16.3). All data are summarized in Table S4.

### Drug resistance and MTBC population structure

1 017 were classified as MDR (54.9 %) and 835 as not MDR, including 681 (36.8 %) pan-susceptible strains and 154 strains with variable resistances other than MDR (8.3 %, Figure 2, Table S3). Among the 1 017 MDR strains, 588 strains were pre-XDR (31.7 % of the total population, and 57.8 % of the MDR MTBC strains), and 182 XDR (9.8 % of the total population, and 17.9 % of the MDR MTBC strains, Figure 2, Table S3, Table S4).

**Figure 2.**
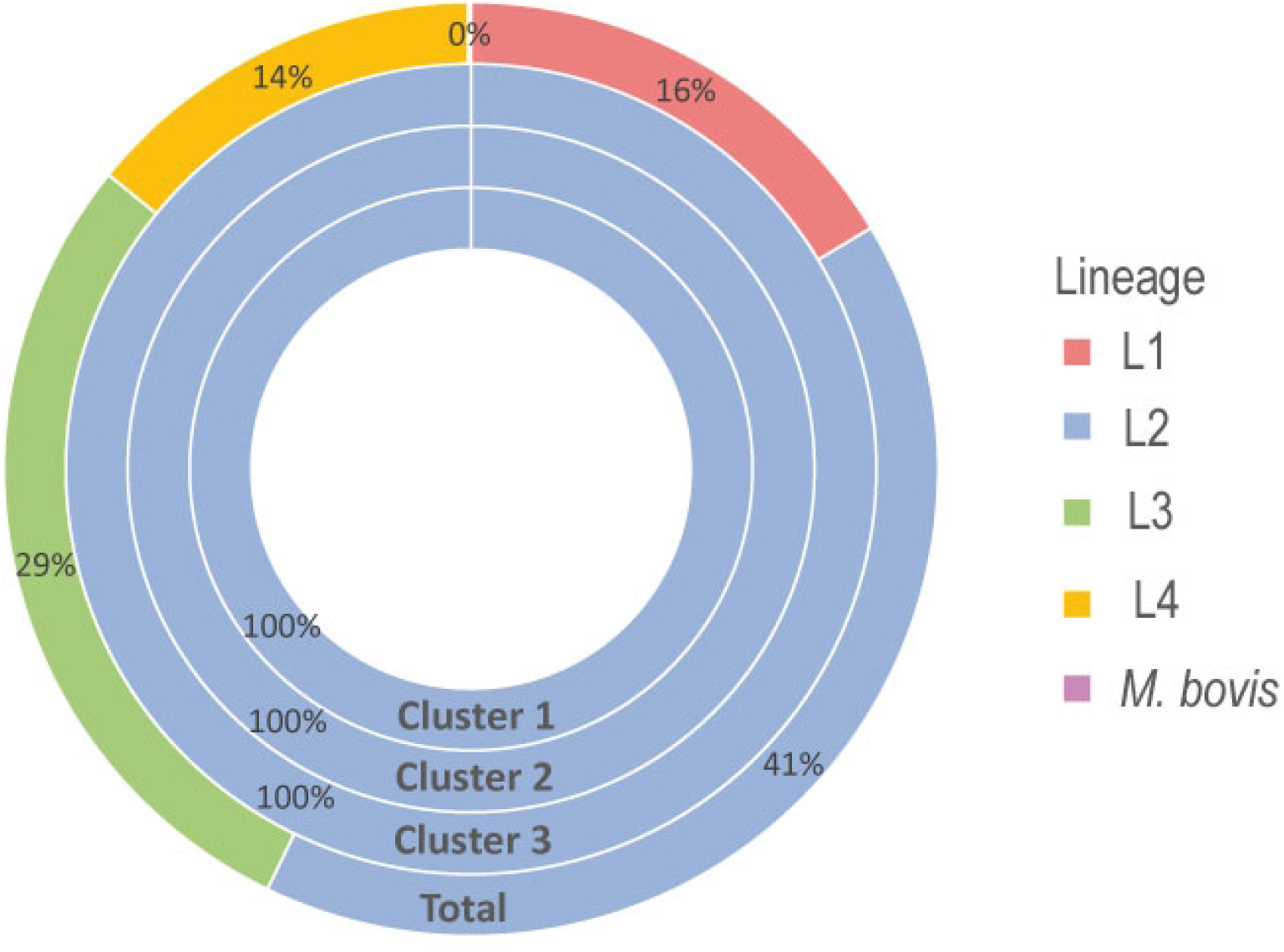
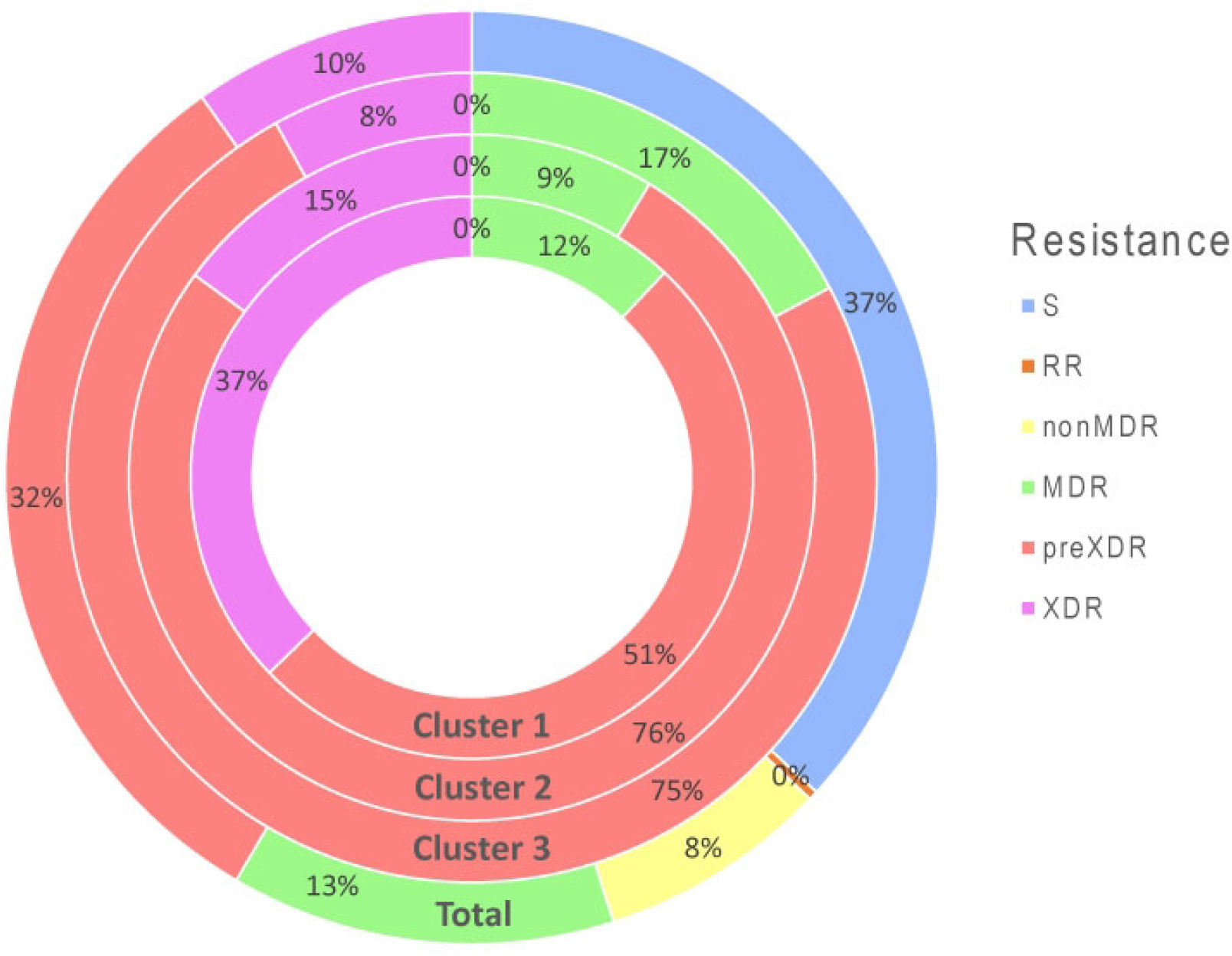
Genotype and Resistance category distribution of strains across 1 852 isolates (Total) and within the three major clusters. **A**. Distribution of genotypes across the 1 852 isolates; Lineage 1 (EAI and EAI Manila), Lineage 2 (Beijing), Lineage 3 (Delhi-CAS) and Lineage 4 (Euro-American, H37Rv-like, Haarlem, LAM, mainly T, S-type, Ural and X-type); For the total dataset the main lineages are Lineage 2 (41 %), followed by Lineage 3 (29 %) and Lineage 1 (16 %). All strains in cluster 1,2 and 3 belong to Lineage 2. **B**. Distribution of resistance category across the 1852 isolates; Resistance categories are RR (Rifampicin Resistant), non MDR (resistant, but not multi drug resistant [MDR]), MDR, pre-XDR (pre-extensively drug resistant) and XDR (extensively drug resistant); of the total strain population about 37 % are susceptible (S) to all drugs, 32 % of strains are pre-XDR whereas 10 % are XDR in the cohort. The resistant category distribution of cluster 1, 2 and 3 differs, with all strains being at least MDR and up to 37 % of strains being already XDR (Cluster 1).

Lineage 2 strains (L2, Beijing/East Asia) constituted 41 % of the total collection (n = 756), followed by Lineage 3 (L3, Delhi/CAS, n = 531, 29 %), Lineage 1 (L1, East Africa Indian, EAI, n = 303, 16 %) and Lineage 4 (L4, Euro-American, n = 260, 14 %) strains (Figure 2). L2 strains are overrepresented in drug resistance strains, especially in those harboring multiple drug resistances, while strains of the other three lineages show an opposite trend. Indeed, 77 % of all pre-XDR and 86 % of all XDR strains belong to L2 (Table S3).

### Genome based cluster analysis

A cgMLST-based cluster analysis employing a threshold of a maximum allele distance of 12 alleles grouped 801 (43 %) of the 1 852 strains into 96 clusters, ranging in size from two to 258 strains (Figure S2, Table S4). The three biggest clusters comprise 258 (cluster 1), 127 (cluster 2) and 87 (cluster 3) strains. Clustered strains comprise all four major MTBC-lineages, however, all strains of the three largest allele clusters were L2 strains (Figure 2, Table S3). If the different resistance categories were considered, the cluster rate was 55.1 % in MDR, 77.7 % in pre-XDR and 84.6 % in XDR strains (Table S3).

Cluster 1-3 strains were close to 100 % resistant to INH, RMP, EMB, PZA, and SM, while only PZA resistance rate was lower in cluster 3 strains (11.5 %, Figure S3) and also developed high FQ resistance rates (84.1 %) rendering them at least pre-XDR (see below). Resistance to injectable drugs was 42.2 % in cluster 1 strains, thus, 96 out of the 258 cluster 1 strains were classified as XDR (Figure 2, Table S3).

### Factors associated with pre-XDR/XDR-TB

In the univariate statistical analysis, sex and age group were not associated with pre-XDR/XDR-TB (Table 1). L2 strains (P < 0.001), belonging to a cluster (P < 0.001), and belonging to clusters 1-3 increased the odds of pre-XDR/XDR compared to MDR, and L1, 3, and 4 strains had a lower odd of being pre-XDR/XDR compared to MDR (Table 1).

**Table 1.** Characteristics of MDR and pre-XDR/XDR cases with factors associated with pre-XDR/XDR-TB based on univariate and multivariate analysis.

|  | MDR |  | pre-XDR/XDR |  | Univariate analysis |  |  |  | Multivariate logistic regression |  |  |  |
| --- | --- | --- | --- | --- | --- | --- | --- | --- | --- | --- | --- | --- |
|  | n | % | n | % | OR | 95% lower | 95% upper | P-value | Adjusted OR | 95% lower | 95% upper | Adjusted P-value |
| Total | 247 | 100% | 770 | 100% |  |  |  |  |  |  |  |  |
| Gender |  |  |  |  |  |  |  |  |  |  |  |  |
| M | 101 | 41% | 333 | 43% | 1.09 | 0.82 | 1.46 | 0.55 |  |  |  |  |
| F | 145 | 59% | 437 | 57% |  |  |  |  |  |  |  |  |
| unknown | 1 | 0% | 0 | 0% |  |  |  |  |  |  |  |  |
| Age |  |  |  |  |  |  |  |  |  |  |  |  |
| <18 | 34 | 14% | 96 | 12% | 0.89 | 0.58 | 1.40 | 0.59 |  |  |  |  |
| 18-40 | 155 | 63% | 502 | 65% | 1.11 | 0.82 | 1.51 | 0.49 |  |  |  |  |
| 40-60 | 41 | 17% | 143 | 19% | 1.15 | 0.77 | 1.72 | 0.51 |  |  |  |  |
| >60 | 16 | 6% | 29 | 4% | 0.57 | 0.29 | 1.14 | 0.08 |  |  |  |  |
| unknown | 1 | 0% | 0 | 0% |  |  |  |  |  |  |  |  |
| Lineage |  |  |  |  |  |  |  |  |  |  |  |  |
| Lineage 1 (EAI) | 37 | 15% | 13 | 2% | 0.10 | 0.05 | 0.19 | < 0.001 | 0.18 | 0.08525 | 0.378876 | < 0.001 |
| Lineage 2 (Beijing) | 103 | 42% | 611 | 79% | 5.36 | 3.90 | 7.39 | < 0.001 | 2.07 | 1.24238 | 3.461374 | 0.01 |
| Lineage 3 (Delhi-CAS) | 69 | 28% | 70 | 9% | 0.26 | 0.18 | 0.38 | < 0.001 | 0.51 | 0.30597 | 0.853136 | 0.01 |
| Lineage 4 (Euro-American) | 38 | 15% | 76 | 10% | 0.60 | 0.39 | 0.94 | 0.02 |  |  |  |  |
| *M. bovis* | 0 | 0% | 0 | 0% |  |  |  |  |  |  |  |  |
| Clustering |  |  |  |  |  |  |  |  |  |  |  |  |
| Yes | 136 | 55% | 611 | 79% | 3.13 | 2.28 | 4.30 | < 0.001 | 0.89 | 0.59700 | 1.31551 | 0.55 |
| No | 111 | 45% | 159 | 21% |  |  |  |  |  |  |  |  |
| Cluster |  |  |  |  |  |  |  |  |  |  |  |  |
| Cluster 1 | 31 | 13% | 227 | 29% | 2.91 | 1.92 | 4.53 | < 0.001 | 1.65 | 0.98703 | 2.754255 | 0.06 |
| Cluster 2 | 11 | 4% | 116 | 15% | 3.80 | 2.00 | 7.97 | 0.003 | 2.37 | 1.16704 | 4.831159 | 0.02 |
| Cluster 3 | 15 | 6% | 72 | 9% | 1.59 | 0.88 | 3.06 | 0.12 | 1.08 | 0.56019 | 2.085229 | 0.82 |

In multivariate analysis, the odds of a strain being pre-XDR/XDR was twice as high for strains belonging to L2 (aOR 2.07, 95% CI 1.24, 3.46) and for strains belonging to a cluster 2 (aOR 2.37, 95% CI 1.17, 4.83), 50% lower for strains belonging to lineage 3 (aOR 0.51, 95% CO 0.31, 0.85) and 80 % lower for strains belonging to lineage 1 (aOR 0.18, 95 % CI 0.31, 0.85; Table 1 and Table S5).

### High resolution SNP-based analysis of cluster 1-3 MTBC strains

To get in-depth data on the transmission dynamics and evolution of the most dominant strains in our collection, we performed a high-resolution SNP-based analysis on the L2 strains of allele clusters 1-3. Using a L2 sub-classification system^8^, cluster 1 strains were classified as Asian/African 2 subgroup, whereas cluster 2 strains were classified as Ancestral 3 strains. Strains from cluster 3 could not be assigned to a particular previously defined L2 subgroup (Table S4).

Using a maximum SNP distance of 12, a total of 239 cluster 1 strains could be grouped into eight SNP-based clusters (SNP_cl), ranging in size from two to 143 isolates (Figure 3, Table S4). The largest SNP cluster is SNP_cl 1 with 143 isolates, followed by SNP_cl 2 and SNP_cl 3, which comprise 37 and 41 isolates, respectively. The phylogeny of the strains in maximum likelihood phylogeny based on the concatenated SNP sequence (356 parsimony-informative, 961 singleton sites, 1 106 constant sites) is in line with particular resistance types, and, thus, confirmed the clonality of the isolates in particular sub-branches e.g. by carrying the same *rpoB, embB*, or *pncA* mutations (Figure 3).

**Figure 3.**
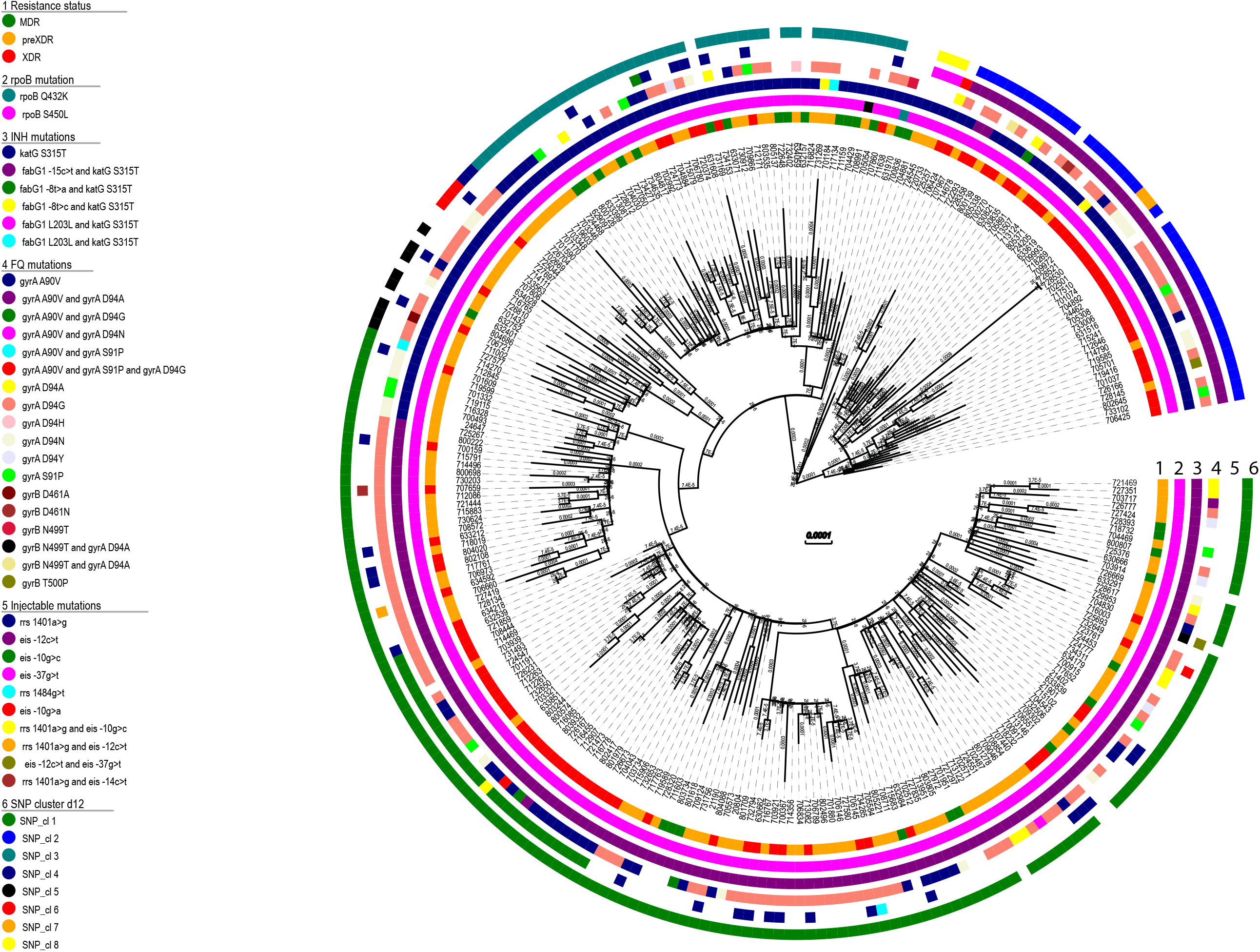
Maximum likelihood phylogeny based on the concatenated SNP sequence of 258 MTBC strains from allele-based cluster 1. The concatenated SNP sequence consists of 356 parsimony-informative, 961 singleton sites and 1 106 constant sites; mutations related to respective drugs and resistance status are color coded and expressed as annotation rings on the tree. SNP-based clusters with maximum distance of 12 (d12) is plotted on the outer ring. Abbreviations: INH, Isoniazid; EMB, Ethambutol; PZA, Pyrazinamide; FQ, Fluoroquinolones.

The close relationship and likely transmission of pre-XDR and XDR strains was further confirmed by the grouping of isolates into several closely related subgroups in the phylogeny that share the same FQ and injectable drug resistance mutations e.g. *eis* -12c/t, *eis* -10g/t, *gyrA* 90V, or even double mutations such as *gyrA* 90V and *gyrA* D94G (Figure 3, Table S4), pointing towards a common ancestor that acquired these mutations before the clone started spreading as pre-XDR/XDR in the Mumbai area. In cluster 1 117 (89.3 %) of the 131 pre-XDR MTBC strains carried a mutation that leads to FQ resistance, which makes those strains already comparable with XDR MTBC strains from a clinical point of view. Only 13 (9.9 %) were resistant to an injectable drug, but susceptible to FQs.

To test if strains of particular subgroups were spreading in certain areas in Mumbai, we linked the SNP clusters with geographical occurrence (Figure 4). However, this analysis confirmed that strains of all cluster 1 SNP subgroups occur in all parts of the study region indicating wide-spread of these clones. Finally, we screened the NGS datasets for compensatory mutations that have been previously described to enhance the transmissibility of MDR strains^8,9,15^. This analysis revealed that 243 of the 258 cluster 1 strains carry at least one compensatory mutation in *rpoC* (Table S4).

**Figure 4:**
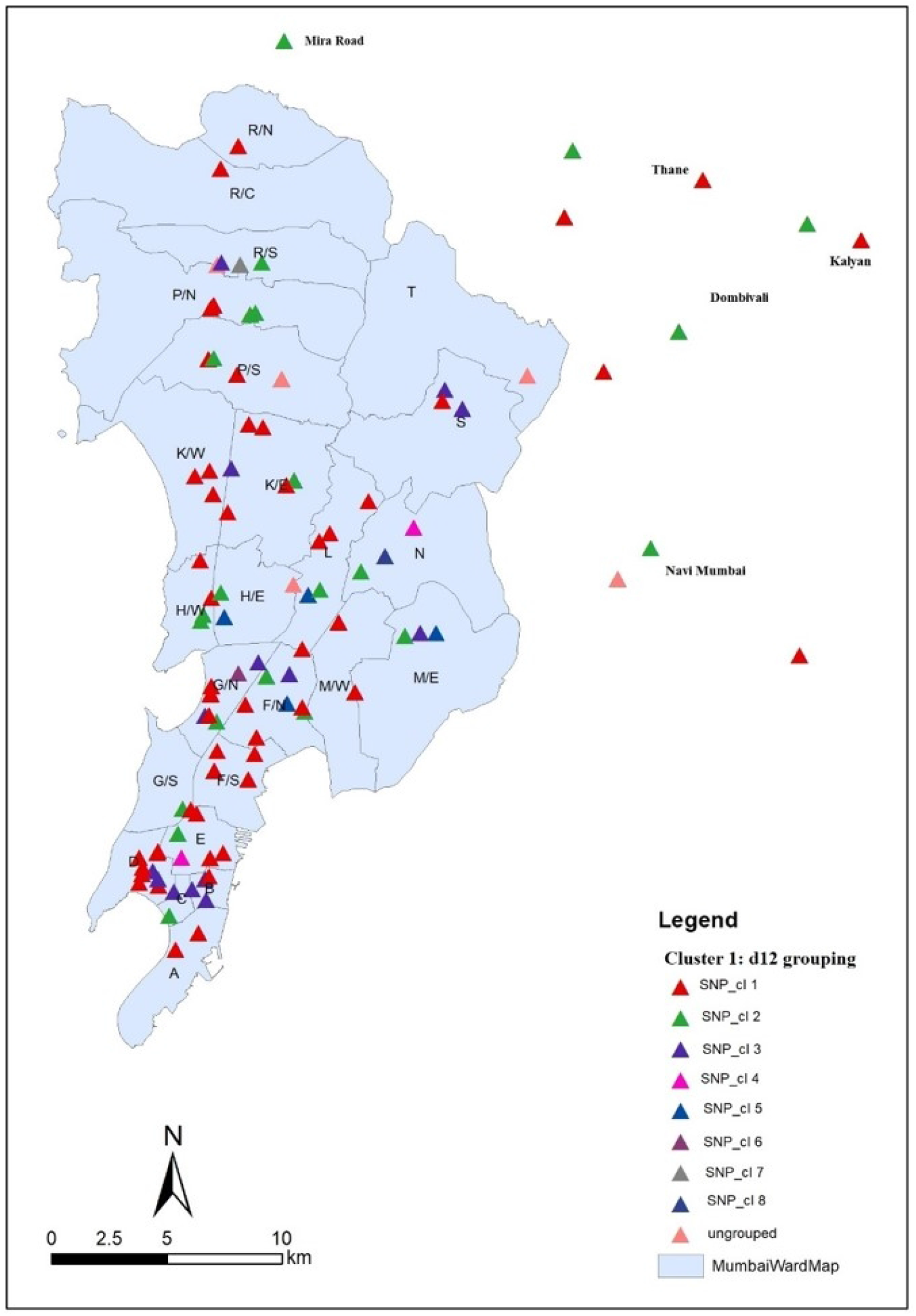
Geographical occurrence of patients with maximum SNPs distance of 12 for Cluster 1 across Mumbai Metropolitan Region (MMR). The geographical distribution underlines the widespread of all cluster 1 SNP subgroups across the city and its neighboring areas. Boundaries of the map for the neighboring regions are not available online.

SNP-based analysis of strains from cluster 2 and 3 confirmed their high clonality with virtually all strains belonging to 1 SNP group (Figure S4). The phylogeny of cluster 2 and 3 strains reveals a similar pattern than observed for cluster 1 strains. Sub-groups share same patterns of resistance mutations indicating their acquisition by a common ancestor followed by clonal transmission (Figure S4). This also applies for pre-XDR and/or XDR MTBC strains.

## Discussion

Our large-scale genome-based study analyzing 1 852 MTBC strains from India, mainly the Mumbai Metropolitan region, revealed several remarkable findings shedding light on the MDR, pre-XDR and XDR epidemic in one of the highest populated metropolitan areas of the world. First, we determined high rates of pre-XDR, and XDR strains among the MDR strain population in this study. Indeed, 58.4 % of all MDR strains are already pre-XDR and 17.9 % are XDR. This remarkable shift towards pre-XDR is mediated by high frequencies of FQ resistance mutations that, in combination with particular injectable drug resistance mutations, result in XDR-TB. In addition, pre-XDR, and XDR strains are virtually all L2 strains and show high cluster rates as an indication of ongoing transmission.

On a global level, the WHO reports that approximately 6 % of the MDR-TB patients have an XDR-TB^2^. With 17.9 %, the XDR-TB rate is clearly higher in our study population. Additionally, we observed a high rate (58.4 %) of pre-XDR strains with FQ resistance, thus, rendering one of the most effective drugs of the short and long MDR-TB regimen non-effective^16^. Indeed, *gyrA* D94G (41%) and *gyrA* A90V (21%) were observed as the predominant mutations in the FQ resistant strains from India in the CRyPTIC project (unpublished data), the former mutation contributing to high level resistance (unpublished data).

To the best of our knowledge, a comparable shift of resistance from MDR towards pre-XDR and XDR has not been reported from other regions, though it has been evident in Mumbai over the last two decades^7,17,18^. Our data underline the potential of MDR MTBC strains to evolve further resistance and yet efficiently transmit even when they become pre-XDR and XDR.

Ongoing and effective transmission of MDR, pre-XDR, and XDR MTBC strains is further confirmed by subgroups of clustered strains, that share identical variant sites conferring resistance including FQ resistance and injectable drug resistance mutations. The clonal expansion of particular pre-XDR/XDR clones with combined resistance to all first line drugs and FQs reduces available group A, B, and C drugs proposed for the treatment of MDR-TB cases to a minimal set. This renders use of the short MDR-TB regimen impossible for patients infected with such strains^2,16^. Geographical mapping showed that strains of the dominant cluster 1 are dispersed across all districts of the Mumbai metropolitan area of more than 28 million inhabitants.

A further striking finding of our study is strong association of L2 strains with pre-XDR/XDR-TB linked with a very high cluster rate (85 %) of L2 strains. Indeed, strains of the three major L2 clusters account for half of the pre-XDR and two thirds of all XDR-TB cases, and just cluster 1 strains account for 52.1 % of the XDR MTBC strains. The proportion of Beijing strains in Mumbai increased over the last two decades^7,17,19^. These data highlight the insights sequence based molecular epidemiological analysis reveals on how an MDR/pre-XDR/XDR epidemic emerges and becomes prevalent in a densely populated geographic area. In fact, the challenge to control pre-XDR/XDR-TB in Mumbai specifically requires intensification of drug resistance detection, effective drug treatment and transmission control of clusters 1-3 strain. Robust monitoring is required for detection and control of new emergent, fit, and transmissible multi-resistant lineages. An earlier study has already highlighted the competitive fitness of MDR strains from India using growth competition experiments ^20^. An association between BCG vaccine escape and efficient spread of Beijing strains has also been proposed^21,22^.

The clonality of the MDR/pre-XDR/XDR MTBC strains in our collection is remarkable. Previously, we determined dominant MDR clones in Eastern European settings, and could show that the spread of a few dominant MDR MTBC strains with compensatory mutations e.g. in *rpoC* can significantly affect the MDR-TB epidemic in specific regions^8,9,23,24^. However, pre-XDR and XDR-TB cases have until now been found at a comparably lower frequency and FQ resistance has not been present in more than 30% of the isolates ^8,9,23,24^. Thus, our data extend these findings to pre-XDR/XDR-TB and demonstrate that pre-XDR/XDR-TB can be propelled by a few lineages that have obviously acquired compensatory mutations, transmit efficiently and reach high prevalence in the population.

Our study has limitations. We only investigated MTBC strains from the laboratory of one tertiary care hospital in Mumbai which represents community samples from a pool of private physicians from the MMR. Although our study covered approx. 16 % of MDR cases in occurring in the region, it may not be representative for the whole Mumbai metropolitan area and for the rest of India. Still, our geographical mapping shows that the study captured cases from the main districts of Mumbai as well as from other areas in India. However, considering our findings, larger investigations on the MDR/pre-XDR/XDR proportions and on the spread of dominant pre-XDR/XDR MTBC strains in Mumbai and other parts of India urgently need to be performed.

In conclusion, our data indicate that the majority of pre-XDR/XDR-TB cases are caused by highly transmissible L2 lineages. Thus, successful control in Mumbai of the DR epidemic urgently requires measures for stopping the transmission of MDR/pre-XDR/XDR strains of the Beijing lineage. As the majority of pre-XDR strains are already FQ resistant, treatment options are limited and rapid adaptation of treatment strategies, for example, comprehensive resistance detection for better design of personalized effective treatment regimens need to be established. It is likely, that the uninformed use of treatment regimens including the newest MDR-TB drugs without precise knowledge of individual resistance patterns and close patient monitoring will likely result in further resistance development as described already^25–28^ and ongoing transmission of even more resistant strains. The national extent of spread in India by the dominant pre-XDR/XDR clones identified in Mumbai and other regions needs to be urgently considered.

## Methods

### Study design

A total of 2 040 MTBC strains from patients were retrospectively collected as random sequential samples for the CRyPTIC Consortium Project between February 2017 and May 2018 (15 months) from the laboratory of a tertiary care hospital in India and sequenced with an WGS approach. Considering that in Mumbai around 5 000 MDR cases are reported annually, the study covered approx. 16 % of MDR cases occurring in the region (n = 1 017/6 250) (source: https://portal.mcgm.gov.in/)^29^. One hundred and eighty-eight isolates were excluded from analysis due to sequence deficiencies. A majority (n = 1 773) of the remaining isolates (n = 1 852) were collected from the Mumbai Metropolitan region (MMR) of the western State of Maharashtra while 46 were derived from distal parts of Maharashtra and neighboring States/Union Territories and 33 from Delek Hospital, Himachal Pradesh, North India. Approval for the CRyPTIC study was obtained from the Health Ministry’s Screening Committee (HMSC), Government of India dated 6^th^ October 2016. Approval was also obtained from the Institutional Ethics Committee (IEC) of The Foundation for Medical Research, Mumbai (Ref nos. FMR/IEC/TB/01a/2015 and FMR/IEC/TB/01b/2015) and Institutional Review Board of P.D. Hinduja Hospital and Medical Research Centre, Mumbai (Ref no. 915-15-CR [MRC]).

### Molecular methods

Genomic DNA was isolated from the 2 040 patient samples using FastPrep24 lysis method (MP Biomedicals, California, USA) as per standard protocol and quantified using Qubit (Life Technologies, Carlsbad, California, USA). Libraries for WGS were prepared using Nextera XT DNA Library Prep Kit and sequencing was performed on the Illumina NextSeq500 machine as per manufacturer’s protocol (Illumina Inc., San Diego, California, USA) producing 2 x 151 base pair reads.

### Genome analysis

All WGS data were analyzed using the MTBSeq pipeline (Version 1.0.3)^30^. Details are described in supplemental methods (Text S1). Phylogenetic lineages (MTBC-lineages and known Beijing subgroups) were inferred from specific SNPs based on Coll *et al*. 2014^31^ and Merker *et al*. 2015^8^.

### Genome based resistance prediction and cluster analysis

Polymorphisms in 27 drug resistance associated genes that are involved in drug resistance mechanisms and three compensatory target genes (*rpoA, rpoC*, compensate fitness effects of *rpoB* mutations in RR strains^15^ and *ahpC* upstream region, compensate fitness effects of catalase [*katG*] deficit in INH resistant strains^32^) were analyzed (Table S2).

Wild type gene sequences (*M. tuberculosis* H37Rv reference sequence or synonymous/silent mutations) were interpreted susceptible, known resistance variants ^33^ were considered as resistant, as well as insertion and deletions in the following genes: *katG, rpoB, pncA, ethA, Rv0678, ald* and *ddn*.

Primary cluster analysis was done using the cgMLST method as described previously ^34^. Reference mapped reads were uploaded to the Ridom SeqSphere+ software (version 7.0.4)^34^ and analyzed with the predefined *M. tuberculosis* cgMLST scheme v2 (version 2.1), comprising 2 891 gene targets. A minimum spanning tree was calculated using a cluster alert of 12 alleles distance and the pairwise ignorance of missing values. The minimum cluster size was set to two.

SNP-based phylogenies were calculated as described previously^33^. The SNP-based phylogenetic analysis was performed by excluding regions annotated as repetitive elements (e.g. PPE and PE-PGRS gene families), InDels, multiple consecutive SNPs in a 12-bp window (possible InDel artefacts or rare recombination scars), and positions in 92 genes implicated in antibiotic resistance (supplementary Table S1). For the phylogenetic reconstruction the SNP frequency was set to 75 % and a genome position was considered valid when 95 % of the combined strains had enough coverage (four reads per direction) at this position.

We used the concatenated sequence alignment to calculate a maximum likelihood phylogeny using IQ-TREE software^35^ with ModelFinder option and ascertainment bias correction. We include also ultrafast bootstrap (UFBoot) approximation with 1 000 replicates combined with a further optimizing step to reduce the risk of overestimating the branch support. Phylogenetic trees were mid-point rooted using FigTree v1.4.4 and annotated using the online tool EvolView^36^.

### Statistics

Descriptive statistics was performed for patients’ demographics as well as for lineages, resistance categories and clustering status of MTBC strains. Data derived from genomic analysis of clinical isolates were analyzed statistically using IBM SPSS Statistics Software for Windows (version 19) and R (version 3.6.1). For univariate analysis of potential factors associated with pre-XDR/XDR TB we performed a Fisher’s exact test. Factors with a significant result in the univariate model were included into a multivariate logistic regression analysis. Odds ratios with 95 % confidence interval (CI) were estimated and variables with P values less than 0.05 were taken as significant characteristics.

## Data availability

Fastq files are available at the European Nucleotide Archive (ENA, Table S1).

## Acknowledgments

Parts of this work have been supported within the CRYPTIC consortium by the Bill & Melinda Gates Foundation [OPP1133541] and the Wellcome Trust [200205/Z/15/Z], the Deutsche Forschungsgemeinschaft (DFG, German Research Foundation) under Germanys Excellence Strategy – EXC 2167 Precision Medicine in Inflammation, and the Leibniz Science Campus Evolutionary Medicine of the LUNG (EvoLUNG). DWC is funded by the UK National Institutes of Health Research (NIHR) Biomedical Research Centre, Oxford. The funders had no role in study design; in the collection, analysis, and interpretation of data; in the writing of the report; and in the decision to submit the paper for publication.

## Author contributions

VD, AM, PD, MM, KK, IB, CU, DWC, NM, and SN conceived the idea and designed the study, and analysed and interpreted the data. All authors contributed to obtaining and assembling the data. VD, AM, and SN wrote the initial draft of the paper. All authors contributed to data interpretation, final draft of the paper and approved the final version of the manuscript.

## Supplemental data

Table S1. ENA accession numbers.

Table S2. Genes analyzed.

Table S3. Characteristics of analyzed samples divided by resistance group.

Table S4. Data table

Table S5. Multivariate logistic regression results

**Figure S1**. Geographical distribution of the strains investigated. Strains were plotted on a map according to the geographical position of the submitting center and color coded by the resistance category. All categories came from the whole study region.

Abbreviations: S-Susceptible to all drugs, nonMDR – resistant but not multi-drug resistant, MDR – multi-drug resistant, pre-XDR – pre-extensively drug resistant, XDR – extensively drug resistant

**Figure S2**. Minimum spanning tree based on the analysis of 2 891 alleles of the core genome of the 1 852 *M. tuberculosis* complex strains investigated. Missing values were ignored for pairwise comparisons. Strains are color-coded by the respective lineage name. EAI and EAI Manila (Lineage 1), Beijing (Lineage 2), Delhi-CAS (Lineage 3) and Euro-American, H37Rv-like, Haarlem, LAM, mainly T, S-type, Ural and X-type (Lineage 4). Allele clusters are highlighted by color-shaded branches.

**Figure S3**. Barplot of resistance profiles of the strains belonging to allele cluster 1, 2 and 3. Resistance profiles in %. A) Resistance profile of allele Cluster 1, B. Resistance profile of allele Cluster 2, C. Resistance profile of allele Cluster 3

Abbreviations: INH -Isoniazid; RMP -Rifampicin; SM -Streptomycin; EMB -Ethambutol; PZA -Pyrazinamide; MFX -Moxifloxacin; LFX -Levofloxacin; CFZ -Clofazimine; KAN -Kanamycin; AMI -Amikacin; CPR -Capreomycin; ETH -Ethionamide; LZD -Linezolid; BDQ -Bedaquiline; CS -Cycloserine; PAS -para-aminosalycilic acid; DEL -Delamanid

**Figure S4**. Maximum likelihood (ML) phylogeny of strains belonging to cluster 2 and 3. Mutations related to respective drugs and resistance status are color coded on the annotation rings of the tree. A. ML phylogenetic tree of the 127 MTBC strains from Cluster 2 phylogeny is based on the concatenated SNP sequence with 1 110 parsimony-informative and 386 singleton sites B. ML phylogenetic tree of the 87 MTBC strains from Cluster 3, phylogeny is based on the concatenated SNP sequence with 122 parsimony-informative and 395 singleton sites Abbreviations: INH -Isoniazid; EMB -Ethambutol; PZA -Pyrazinamide; FQ -Fluoroquinolones.

