## Supplemental Text and Figures for "Transmission of pre-XDR and XDR-TB in the Mumbai Metropolitan Region, India" (from another version of this preprint: version 1, 2021-02-03)

### Appendix

**Table of contents**


1. **Text:**

   Supplemental methods

   CRyPTIC publication members

2. **Figures**

   Figure S1. Geographical distribution of study population.
   Figure S2. Minimum spanning tree based on cgMLST.
   Figure S3. Resistance profiles of cluster 1-3.
   Figure S4. Maximum likelihood phylogenies of cluster 2 and 3.

3. **Tables**

   Table S1. ENA accession numbers.
   Table S2. Genes analyzed.
   Table S3. Characteristics of analyzed samples divided by resistance group.
   Table S4. Data table
   Table S5. Multivariate logistic regression results

**Appendix 1: Methods**

**Genome analysis**

The reads were mapped to the reference sequence *M. tuberculosis* H37Rv (GenBank ID: NC_000962.3) with the Burrows-Wheeler Aligner (BWA)[1]. Initial mapping was refined using tools from the Genome Analysis Toolkit (GATK)[2] and SAMtools[3] to e.g. exclude PCR artefacts, correct alignment errors around small insertions/deletions (InDels) and recalibrate base quality scores. Minimum criteria for variants (SNPs and InDels) were set to four reads coverage per direction (forward and reverse) and a variant frequency of 20 % for resistance prediction and 75 % for phylogenetic analysis, respectively.

**Genome based cluster analysis**

Cluster analysis was done using the cgMLST method as described previously [4]. Reference mapped reads were uploaded to the Ridom SeqSphere+ software (version 7.0.4)[4] and analyzed with the predefined *M. tuberculosis* cgMLST scheme v2 (version 2.1), comprising 2 891 gene targets. A minimum spanning tree was calculated using a cluster alert of 12 alleles distance and the pairwise ignorance of missing values. The minimum cluster size was set to two.

The SNP-based phylogenetic analysis was performed by excluding regions annotated as repetitive elements (e.g. PPE and PE-PGRS gene families), InDels, multiple consecutive SNPs in a 12-bp window (possible InDel artefacts or rare recombination scars), and positions in 92 genes implicated in antibiotic resistance (supplementary Table S1). For the phylogenetic reconstruction the SNP frequency was set to 75 % and a genome position was considered valid when 95 % of the combined strains had enough coverage (four reads per direction) at this position.

We used the concatenated sequence alignment to calculate a maximum likelihood phylogeny using IQ-TREE software[5] with ModelFinder option and ascertainment bias correction. We include also ultrafast bootstrap (UFBoot) approximation with 1000 replicates combined with a further optimizing step to reduce the risk of overestimating the branch support. Phylogenetic trees were mid-point rooted using FigTree v1.4.4 and annotated using the online tool EvolView[6].

**Statistics**

Descriptive statistics was performed for patients' demographics as well as for lineages, resistance categories and clustering status of MTBC strains. Data derived from genomic analysis of clinical isolates were analyzed statistically using IBM SPSS Statistics Software for Windows (version 19) and R (version 3.6.1). For univariate analysis of potential factors associated with pre-XDR/XDR TB we performed a Fisher's exact test. Factors with a significant result in the univariate model were included into a multivariate logistic regression analysis. Odds ratios with 95 % confidence interval (CI) were estimated and variables with P values less than 0.05 were taken as significant characteristics.

**CRyPTIC publication members list**

Derrick W Crook, Timothy EA Peto, A Sarah Walker, Sarah J Hoosdally, Ana L Gibertoni Cruz, Joshua Carter, Sarah Earle, Samaneh Kouchaki, Yang Yang, Timothy M Walker, Philip W Fowler, Daniel Wilson and David A Clifton, University of Oxford;

Zamin Iqbal, Martin Hunt and Jeff Knaggs, European Bioinformatics Institute;

Daniela M Cirillo, Emanuele Borroni, Simone Battaglia, Arash Ghodousi, Andrea Spitaleri and Andrea Cabibbe, Emerging Bacterial Pathogens Unit, IRCCS San Raffaele Scientific Institute, Milan;

Sabira Tahseen, National Tuberculosis Control Program Pakistan, Islamabad; Kayzad Nilgiriwala and Sanchi Shah, The Foundation for Medical Research, Mumbai;

Camilla Rodrigues, Priti Kambli, Utkarsha Surve and Rukhsar Khot, P.D. Hinduja National Hospital and Medical Research Centre, Mumbai;

Stefan Niemann, Thomas Kohl and Matthias Merker, Research Center Borstel; Harald Hoffmann, Katharina Todt and Sara Plesnik, Institute of Microbiology & Laboratory Medicine, IML red, Gauting;

Nazir Ismail, Shaheed Vally Omar, Lavania Joseph Dumisani Ngcamu, Nana Okozi and Shen Yuan Yao, National Institute for Communicable Diseases, Johannesburg;

Guy Thwaites, Thuong Nguyen Thuy Thuong, Nhung Hoang Ngoc and Vijay Srinivasan, Oxford University Clinical Research Unit, Ho Chi Minh City;

David Moore, Jorge Coronel and Walter Solano, London School of Hygiene and Tropical Medicine and Universidad Peruana Cayetano Heredia, Lima;

George F Gao, Guangxue He, Yanlin Zhao, Aijing Ma and Chunfa Liu, China CDC, Beijing;

Baoli Zhu, Institute of Microbiology, CAS, Beijing;

Ian Laurenson and Pauline Claxton, Scottish Mycobacteria Reference Laboratory, Edinburgh;

Robert J Wilkinson, University of Cape Town, Imperial College London and Francis Crick Institute;

Anastasia Koch, University of Cape Town;

Ajit Lalvani, Imperial College London;

James Posey, CDC Atlanta;

Jennifer Gardy, University of British Columbia;

Jim Werngren, Public Health Agency of Sweden;

Nicholas Paton, National University of Singapore;

Ruwen Jou, Mei-Hua Wu, Yu-Xin Xiao, CDC Taiwan;

Lucilaine Ferrazoli, Rosangela Siqueira de Oliveira, Institute Adolfo Lutz, São Paulo.

James Millard, Africa Health Research Institute, Durban

Rob Warren - University of Stellenbosch, Cape town

Annelies Van Rie, University of Antwerp

Simon Grandjean Lapierre, Marie-Sylvianne Rabodoarivelo and Niaina Rakotosamimanana, Institut Pasteur de Madagascar

Camus Nimmo, University College London

Kimberlee Musser and Vincent Escuyer, Wadsworth Center, New York

Ted Cohen, Yale University

Figure S1

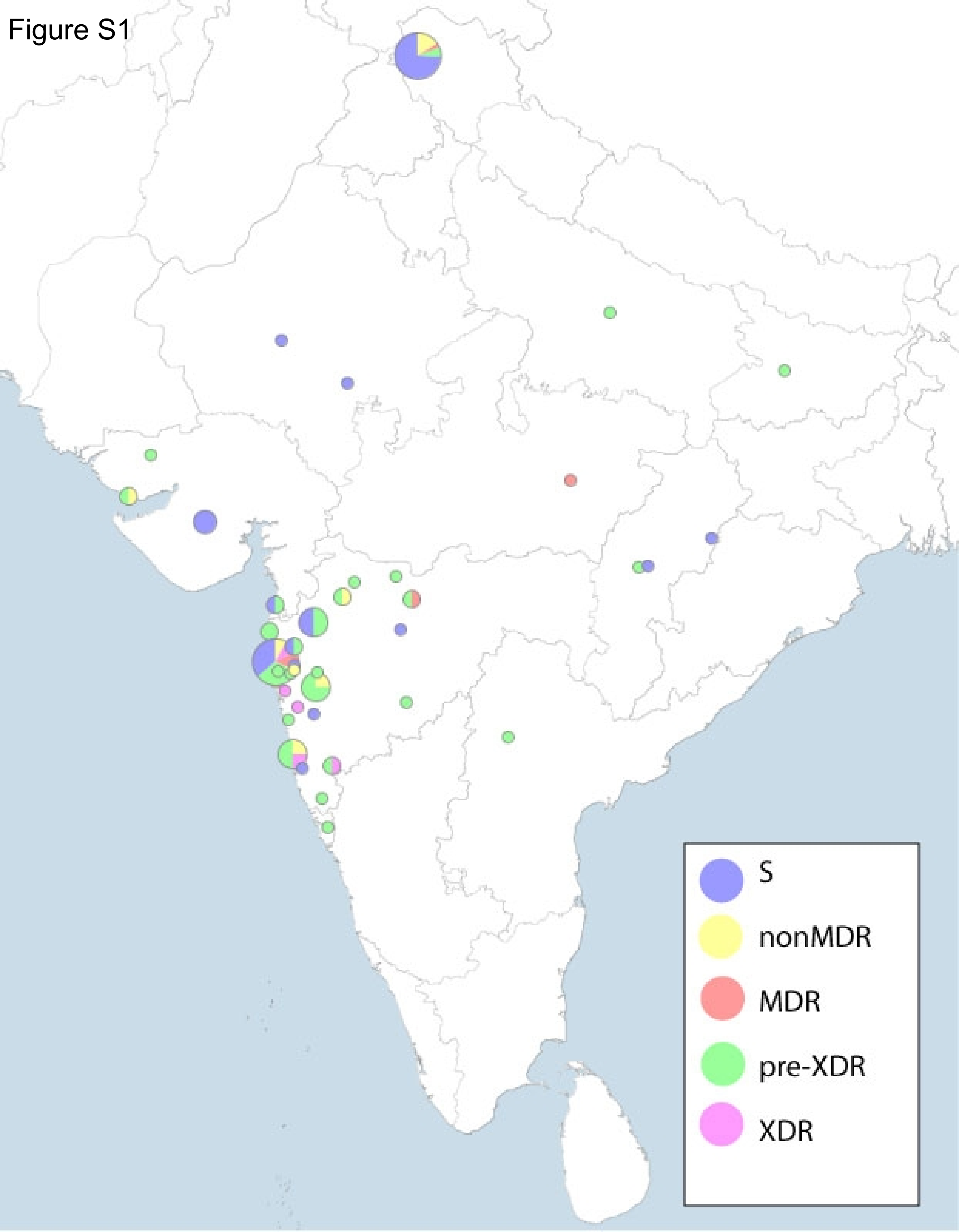

Figure S2

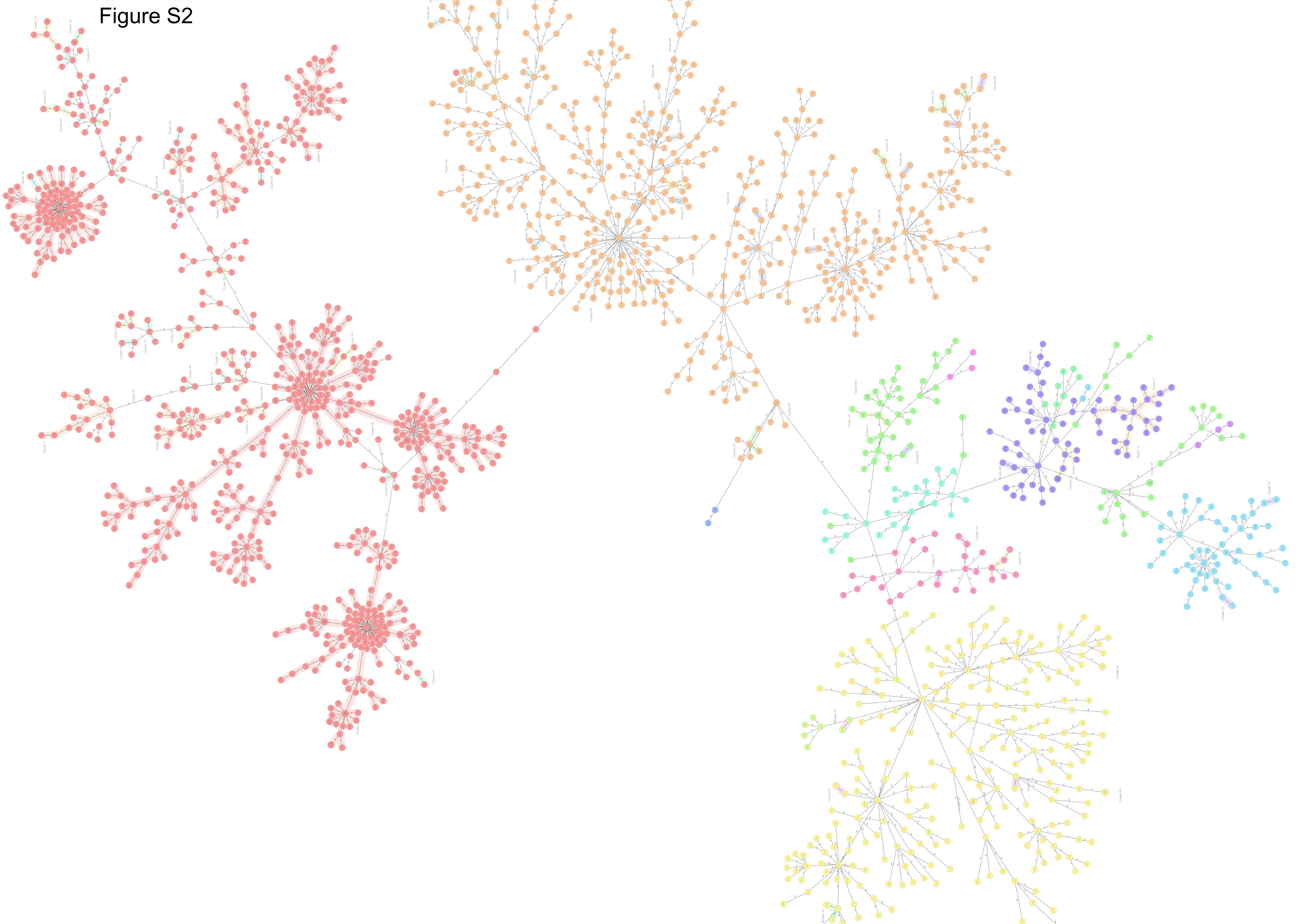

Task Templates: M. tuberculosis complex cgMLST v2 v2.1

Comparison Table Retrieval: CRyPTIC_india [unstored]

Projects: CRyPTIC_india (Mycobacterium tuberculosis complex/bovis/africanum/canetti)

Comparison Table created: Oct 16, 2020 8:23 AM (v7.0.4_(2020-05))

M. tuberculosis complex cgMLST v2 Complex Type / Cluster-Alert distance: 12

Ridom SeqSphere+ MST for 1852 Samples based on 2891 columns, pairwise ignoring missing values

Distance based on columns from M. tuberculosis complex cgMLST v2:cgMLST (2891)

For citing correctly in publications the tools used for this analysis see menu Help | Citations and Licenses.

Cluster distance threshold: 12

Color grouped by column "Borstel lineage":

- Beijing
- Delhi-CAS
- EAI
- EAI Manila
- Euro-American
- H37Rv-like
- Haarlem
- LAM
- M. bovis
- mainly T
- S-type
- Ural
- X-type

Figure S3 A

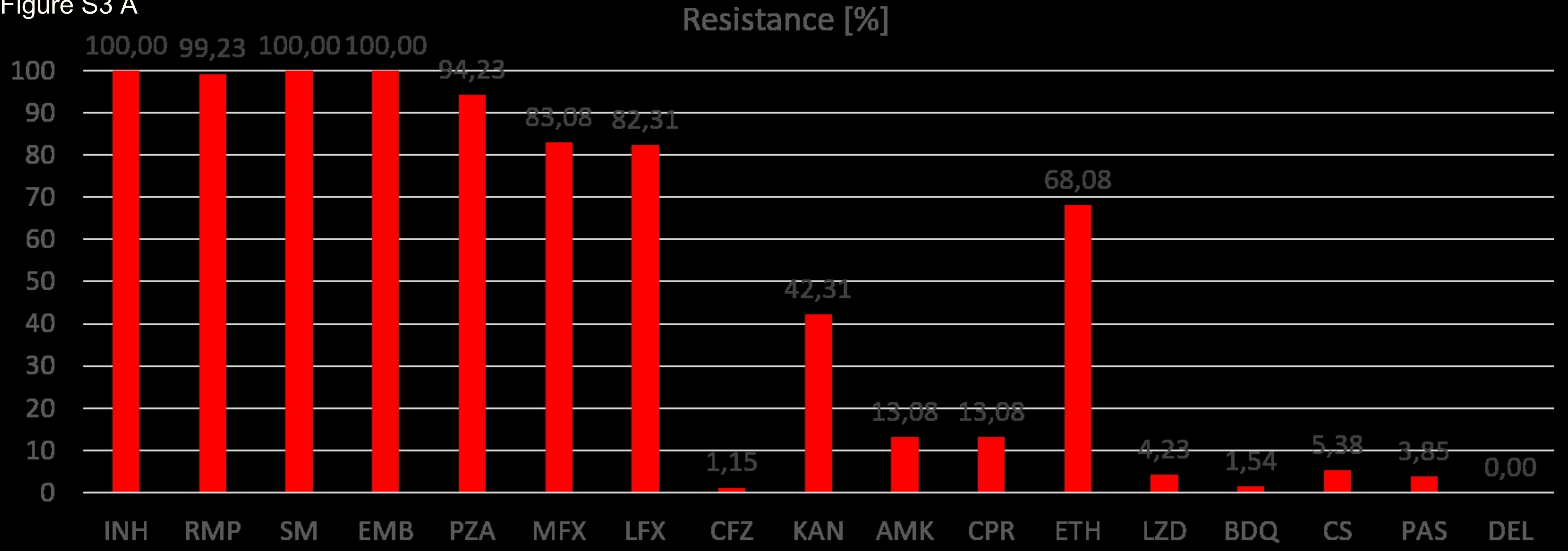

Resistance [%]
100,00
99,23
100,00
100,00
94,23
83,08
82,31
1,15
42,31
13,08
13,08
68,08
4,23
1,54
5,38
3,85
0,00
100
90
80
70
60
50
40
30
20
10
0
INH
RMP
SM
EMB
PZA
MFX
LFX
CFZ
KAN
AMK
CPR
ETH
LZD
BDQ
CS
PAS
DEL

Figure S3 B

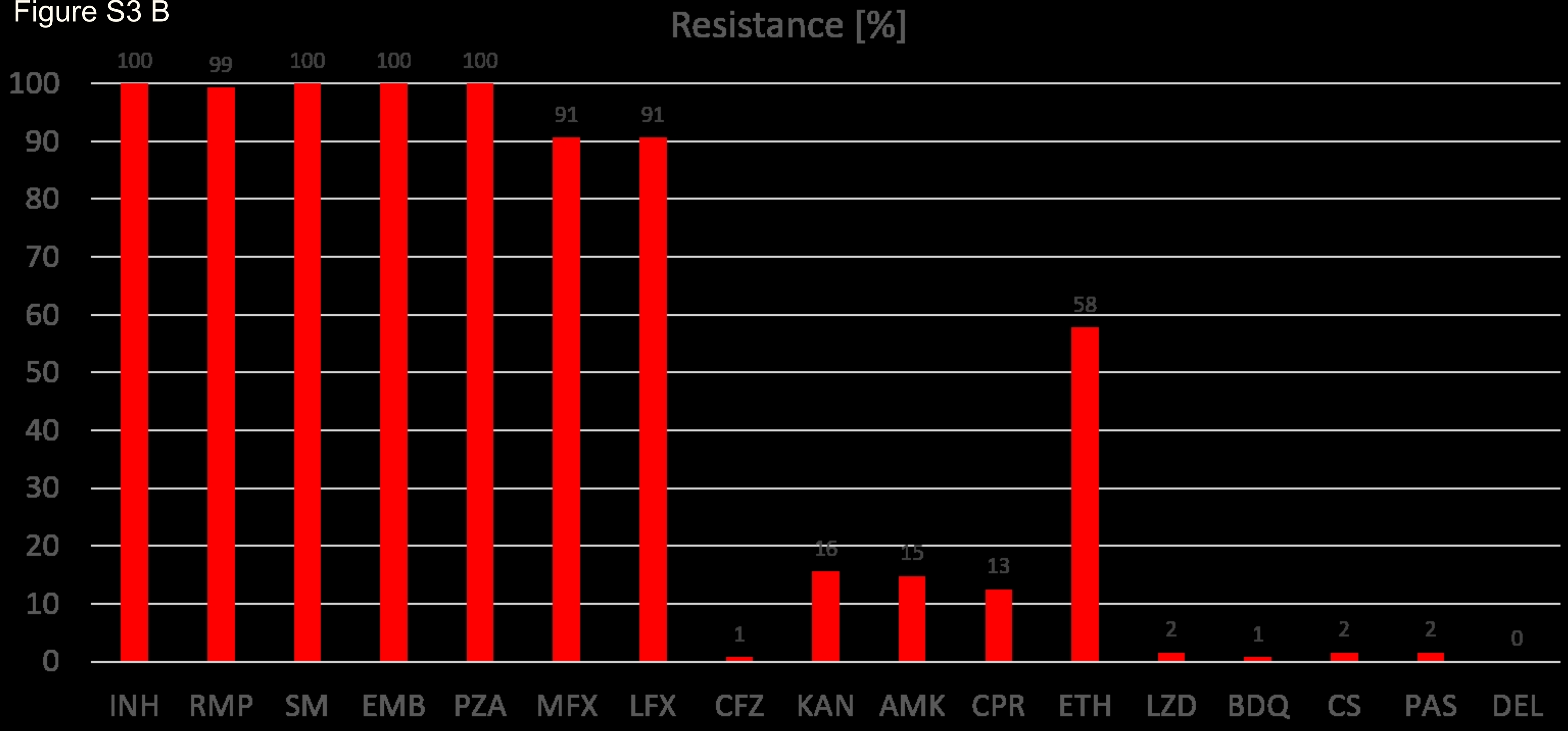

Resistance [%]
100
99
100
100
100
91
91
1
16
15
13
58
2
1
2
2
0
100
90
80
70
60
50
40
30
20
10
0
INH
RMP
SM
EMB
PZA
MFX
LFX
CFZ
KAN
AMK
CPR
ETH
LZD
BDQ
CS
PAS
DEL

Figure S3 C

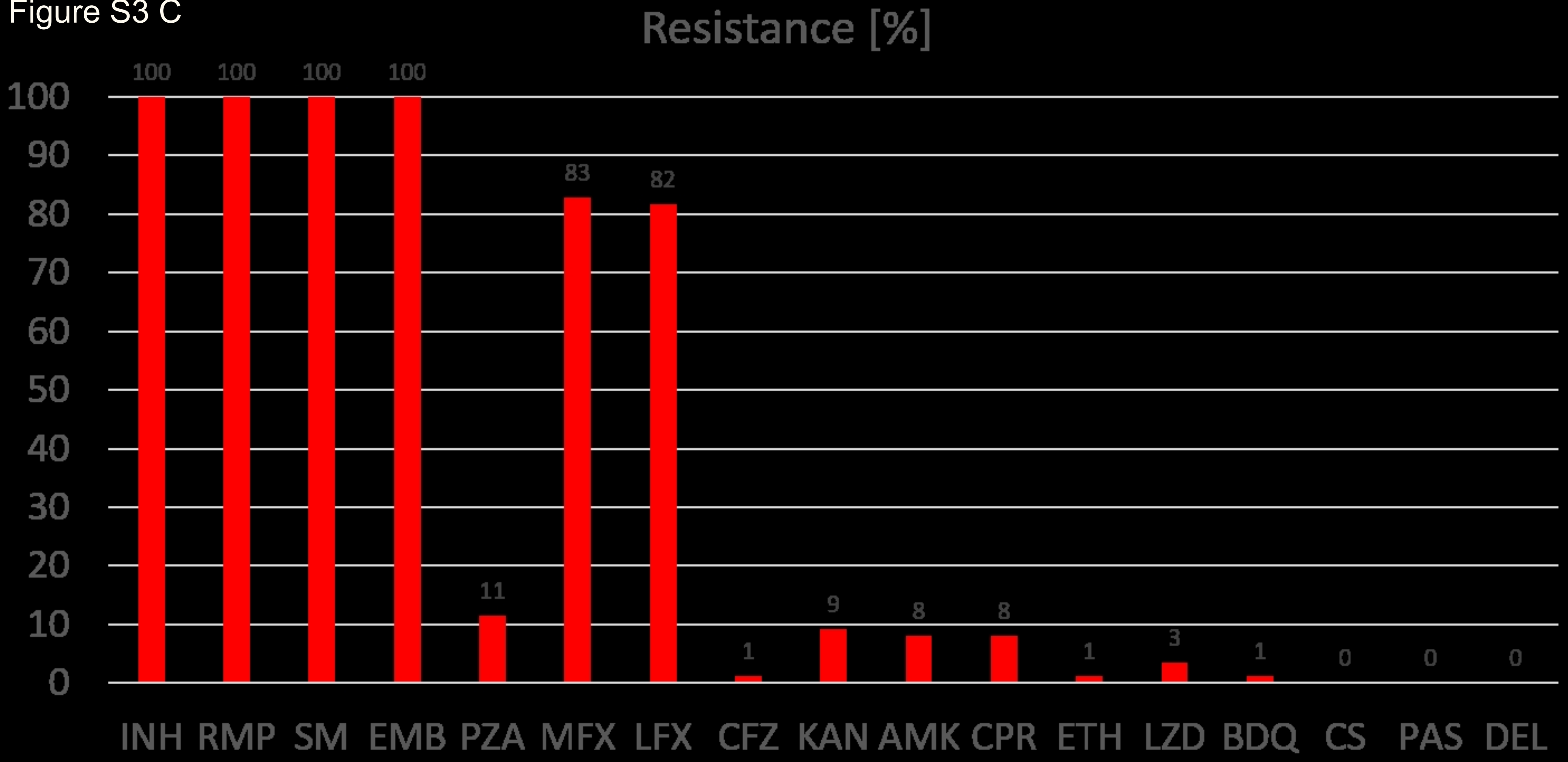
Resistance [%]
100
90
80
70
60
50
40
30
20
10
0
100
100
100
100
11
83
82
1
9
8
8
1
3
1
0
0
0
INH
RMP
SM
EMB
PZA
MFX
LFX
CFZ
KAN
AMK
CPR
ETH
LZD
BDQ
CS
PAS
DEL

### Figure S4 A

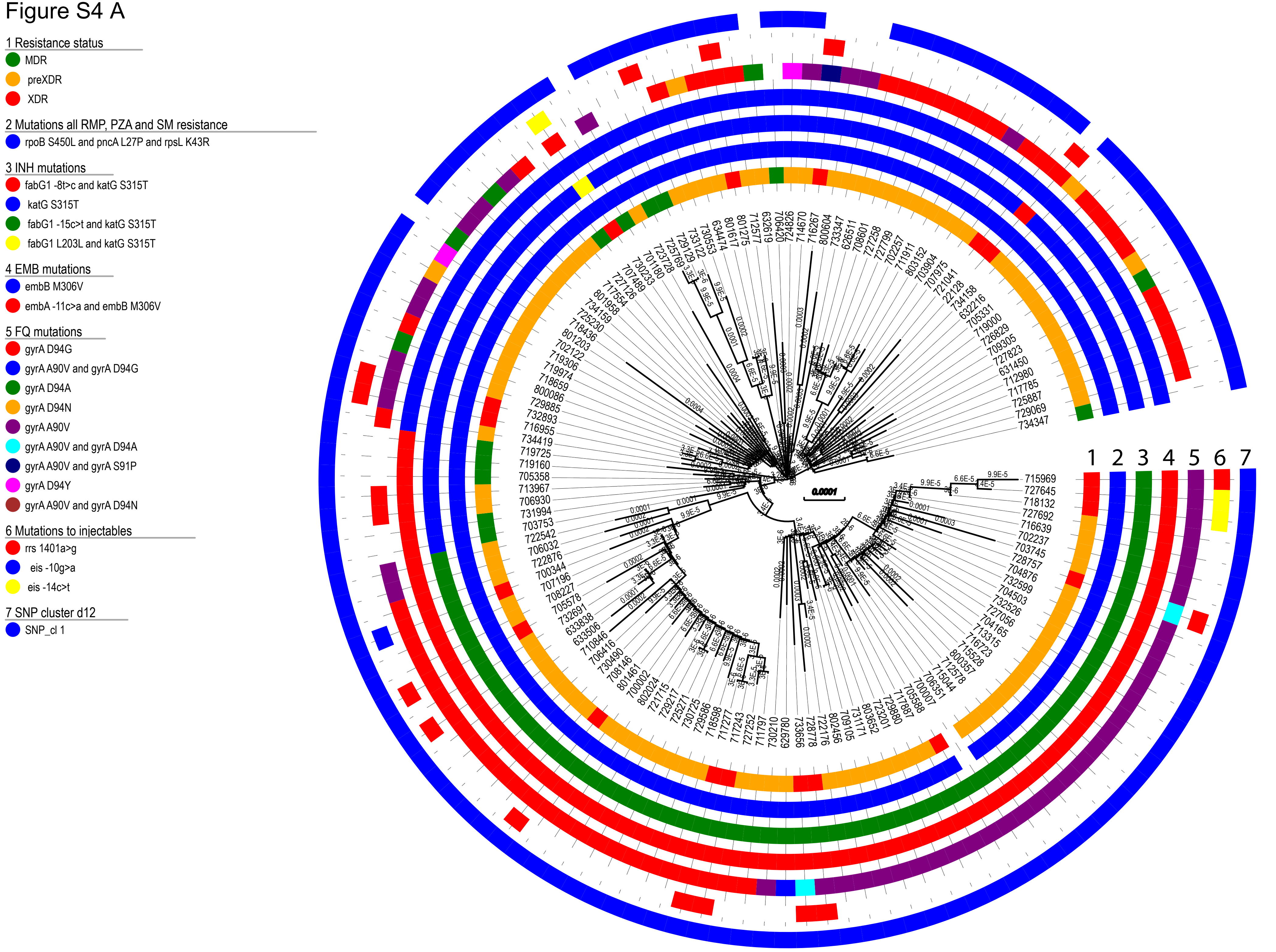
1 Resistance status
MDR
preXDR
XDR
2 Mutations all RMP, PZA and SM resistance
rpoB S450L and pncA L27P and rpsL K43R
3 INH mutations
fabG1 -8t>c and katG S315T
katG S315T
fabG1 -15c>t and katG S315T
fabG1 L203L and katG S315T
4 EMB mutations
embB M306V
embA -11c>a and embB M306V
5 FQ mutations
gyrA D94G
gyrA A90V and gyrA D94G
gyrA D94A
gyrA D94N
gyrA A90V
gyrA A90V and gyrA D94A
gyrA A90V and gyrA S91P
gyrA D94Y
gyrA A90V and gyrA D94N
6 Mutations to injectables
rrs 1401a>g
eis -10g>a
eis -14c>t
7 SNP cluster d12
SNP_cl 1
1 2 3 4 5 6 7
0.0001

Figure S4 B

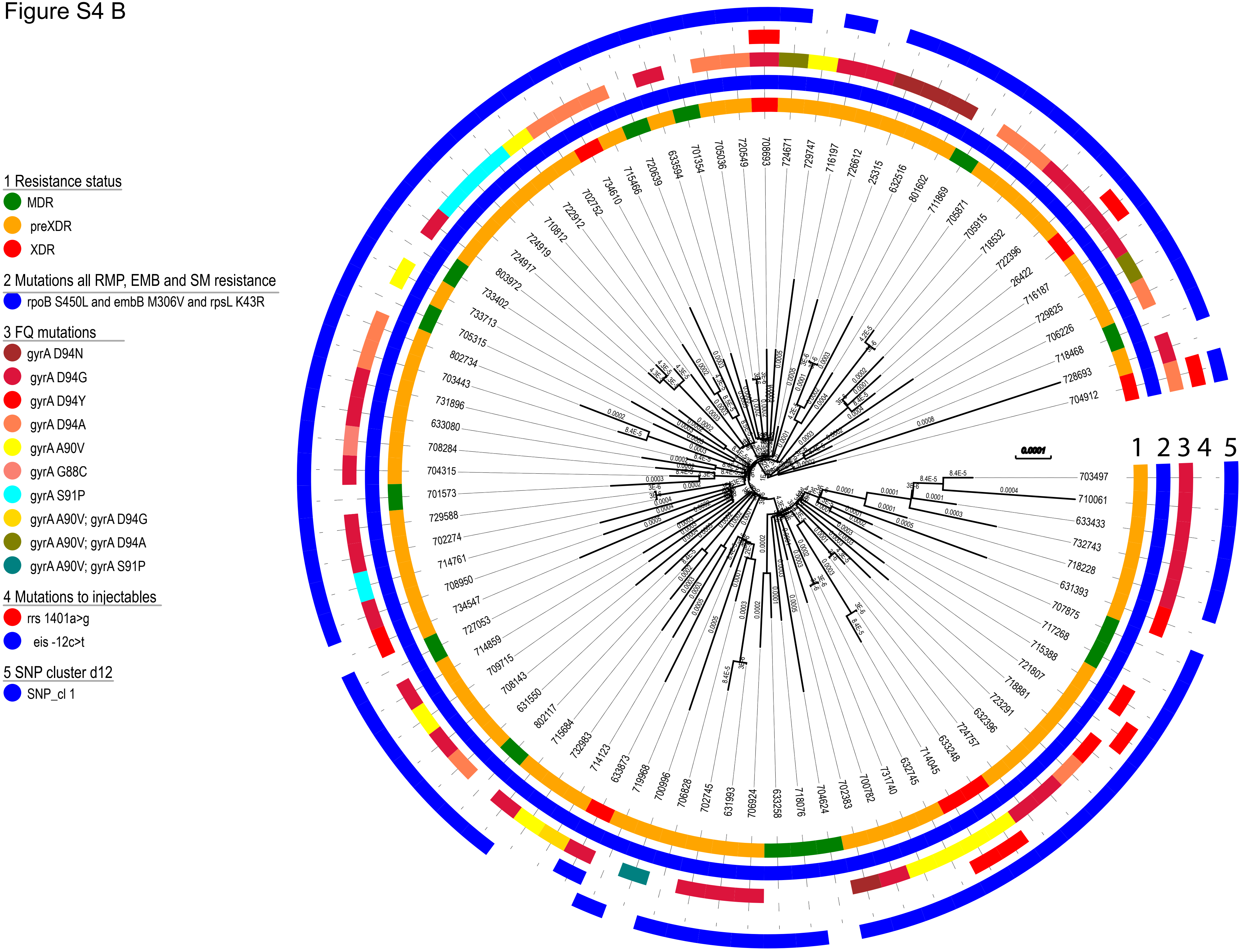

1 Resistance status
MDR
preXDR
XDR
2 Mutations all RMP, EMB and SM resistance
rpoB S450L and embB M306V and rpsL K43R
3 FQ mutations
gyrA D94N
gyrA D94G
gyrA D94Y
gyrA D94A
gyrA A90V
gyrA G88C
gyrA S91P
gyrA A90V; gyrA D94G
gyrA A90V; gyrA D94A
gyrA A90V; gyrA S91P
4 Mutations to injectables
rrs 1401a>g
eis -12c>t
5 SNP cluster d12
SNP_cl 1
1 2 3 4 5
0.0001
708693
724671
729747
716197
726612
25315
632516
801602
711869
705871
705915
718532
722396
26422
716187
729825
706226
718468
728693
704912
703497
710061
633433
732743
718228
631393
707875
717268
715388
721807
718881
723291
632396
724757
633248
714045
632745
731740
700782
702383
704624
718076
633258
706924
631993
702745
706828
700996
719968
633873
714123
732983
715684
802117
631550
708143
709715
714859
727053
734547
708950
714761
702274
729588
701573
704315
708284
633080
731896
703443
802734
705315
733713
733402
803972
724917
724919
710812
722912
702752
734610
715466
720639
633594
701354
705036
720549
